# Epigenomic, transcriptomic and proteomic characterizations of reference samples

**DOI:** 10.1101/2024.09.09.612110

**Authors:** Chirag Nepal, Wanqiu Chen, Zhong Chen, John A. Wrobel, Ling Xie, Wenjing Liao, Chunlin Xiao, Adrew Farmer, Malcolm Moos, Wendell Jones, Xian Chen, Charles Wang

## Abstract

A variety of newly developed next-generation sequencing technologies are making their way rapidly into the research and clinical applications, for which accuracy and cross-lab reproducibility are critical, and reference standards are much needed. Our previous multicenter studies under the SEQC-2 umbrella using a breast cancer cell line with paired B-cell line have produced a large amount of different genomic data including whole genome sequencing (Illumina, PacBio, Nanopore), HiC, and scRNA-seq with detailed analyses on somatic mutations, single-nucleotide variations (SNVs), and structural variations (SVs). However, there is still a lack of well-characterized reference materials which include epigenomic and proteomic data. Here we further performed ATAC-seq, Methyl-seq, RNA-seq, and proteomic analyses and provided a comprehensive catalog of the epigenomic landscape, which overlapped with the transcriptomes and proteomes for the two cell lines. We identified >7,700 peptide isoforms, where the majority (95%) of the genes had a single peptide isoform. Protein expression of the transcripts overlapping CGIs were much higher than the protein expression of the non-CGI transcripts in both cell lines. We further demonstrated the evidence that certain SNVs were incorporated into mutated peptides. We observed that open chromatin regions had low methylation which were largely regulated by CG density, where CG-rich regions had more accessible chromatin, low methylation, and higher gene and protein expression. The CG-poor regions had higher repressive epigenetic regulations (higher DNA methylation) and less open chromatin, resulting in a cell line specific methylation and gene expression patterns. Our studies provide well-defined reference materials consisting of two cell lines with genomic, epigenomic, transcriptomic, scRNA-seq and proteomic characterizations which can serve as standards for validating and benchmarking not only on various omics assays, but also on bioinformatics methods. It will be a valuable resource for both research and clinical communities.

## Introduction

Based on a multi-center whole-genome and whole-exome sequencing study, the FDA Sequencing Quality Control (SEQC-2) consortium has created a set of reference variant datasets derived from the paired cell lines HCC1395 and HCC1395BL and provided best practice guidelines for data analysis^1,2,3^. These two cell lines were from a 43-year-old, white female patient with ductal carcinoma. HCC1395 is an epithelial cancer cell line derived from the mammary gland, while HCC1395BL was prepared from a normal B lymphoblast isolated from the peripheral blood of the same patient^4^. The HCC1395 and its paired normal cell HCC1395BL have been used extensively in breast cancer pathobiology and continue to provide rich information for breast cancer gene discovery, gene expression pathways, and drug discovery^5^. The SEQC-2 consortium also reported comparative analyses of variant calling on whole genome and whole exome data sets^1,2,3^, genomic instability and somatic structural variants (SV)^6^ and showed personalized genome reference significantly improved the detection accuracy of somatic SNVs and SVs^7^. We also evaluated the gene expression profiles at single-cell resolution for these two paired cell lines through a multi-center study and compared their consistency across different centers and different single-cell RNA sequencing (scRNA-seq) technologies^8^. The unique reference resources and methodology established in these studies will notably facilitate precision oncology analysis and personalized cancer therapy.

While the above studies focused on genomic alterations, there is little information regarding the epigenomic and proteomic landscapes of these two reference cell lines, limiting their utility as reference materials. Unlike genomic alterations, epigenetic alterations are frequently reversible chemical modifications that alter DNA accessibility and eventual gene transcription. DNA methylation^9^, chromatin accessibility^10^, and histone modifications^11^ are commonly used to measure epigenetic modifications of DNA within cells. The assay for transposase-accessible chromatin with sequencing (ATAC-seq) measures DNA accessibility and identifies open chromatin regions^12^, which are generally indicative of active transcription. DNA methylation is a biological process by which methyl groups are added to the DNA molecule and is generally a repressive mark. Various protocols like Methyl-seq, Reduced Representation Bisulfite Sequencing (RRBS) or whole genome bisulfite sequencing (WGBS) are used to measure DNA methylation levels^13^, which are frequently reported as bulk methylation percentages defined as beta values (ranging between 0- 1). Proteins, including phosphoproteins^14^, can be identified and quantified by mass spectrometry-based methods. These analyses allow detection of altered protein expression levels or functions through somatic mutations and/or post-translational modifications. Before gene transcription, promoters undergo remodeling of epigenetic marks (e.g., removal of repressive DNA methylation) and open chromatin for transcription factors to bind around transcription start sites. Thus, it is important to understand the nature and extent of epigenetic remodeling, how it influences gene expression, and eventually protein expression and function. Furthermore, we intent to determine the degree of variation in gene expression across cell types that can be explained by epigenetic modifications. To get a comprehensive picture of epigenomic landscapes, we need to measure chromatin accessibility and methylation status simultaneously to understand their intricate relationship in the context of genomic sequence.

Here, we aim to provide a high-quality reference epigenomic, transcriptomic and proteomic map of two cell lines studied by SEQC-2. To this end, we performed ATAC-seq, Illumina TruSeq Capture Epic Methyl-seq, RNA-seq, and proteomic analyses of the HCC1395 and HCC1395BL cell lines and provided a comprehensive catalog of the epigenomic landscape (open chromatin regions and methylation status), transcriptomic and proteomic quantifications. With the detailed annotation of molecular (from genome, epigenome to transcriptome and proteome) snapshot from cells, we sought to understand the inter-relationships across different molecular layers of omic data. We observed that open chromatin regions had low methylation, while closed chromatin regions had high methylation, reflecting a consistent relationship between two epigenetic modifications. This intricate relationship largely coincides with genomic CG density, where CG-rich regions had more accessible chromatin, low methylation levels, and higher gene and protein expression levels. The CG-poor regions had largely repressive epigenetic marks (less open chromatin and higher DNA methylation), often resulting in a cell line or tissue specific expression. Our work provides cell-line reference materials which are well-characterized across epigenomic, transcriptomic and proteomic levels, and highlights the relationship of epigenetic modification with genomic CG context as well as related protein expression.

## Results

### Construction of a reference epigenomic, transcriptomics and proteomic landscapes for HCC1395 and HCC1395BL cell lines

The HCC1395 and HCC1395BL cell lines have been extensively studied including whole-genome and whole-exome sequencing^1–3^, personal genome assembly analysis^7^, scRNA-seq for benchmarking bioinformatics algorithms^8,15^, and structural variation analysis^6^. Currently, these cell lines lack epigenetics and proteomic landscapes, thus we sought to fill this gap by providing a high-quality reference dataset. These two cell lines are from the same donor where HCC1395BL is a cell line derived from a normal B lymphocyte and HCC1395 is a cell line derived from a breast cancer (**Fig. 1**). To measure the epigenetic landscape, we performed two assays, ATAC-seq (to measure chromatin accessibility) and TruSeq Capture Epic Methyl-seq (to measure DNA methylation levels). We used Illumina-based sequencing and performed ATAC-seq on three replicates of HCC1395 (average: 33.14 million (M) paired-end (PE) raw reads) and HCC1395BL (average: 50.56M PE reads) (**Fig. 1**). Similarly, we performed three methyl-seq replicates on HCC1395 (average: 33.14M PE reads) and HCC1395BL (average: 50.56M PE reads) (**Supplementary Table S1**). To understand the association between epigenomics (chromatin accessibility and DNA methylation) with gene expression, we performed three replicates of bulk RNA-seq on HCC1395 (average: 62.6M PE reads) and HCC1395BL (average: 64.04M PE reads). We also integrated scRNA-seq data of these two cell lines from our previous study^8^. In addition, we performed mass spectrometry on three replicates each of HCC1395 and HCC1395BL, resulting in identification and quantification of thousands of expressed proteins. Each data type was systematically analyzed through standard pipelines (**Fig. 1**), such as MACS^16^, Bismark^17^, Kalisto^18^, Seurat^19^ and Preseus^20^ (Methods). After annotation of ATAC-seq peaks and (un)methylated regions, we sought to interrogate the influence of epigenetic patterns on gene and protein expression as well as understand how the genomic DNA (CG-rich or CG-poor) of promoters influenced epigenetic markers and gene expression.

**Figure 1.**
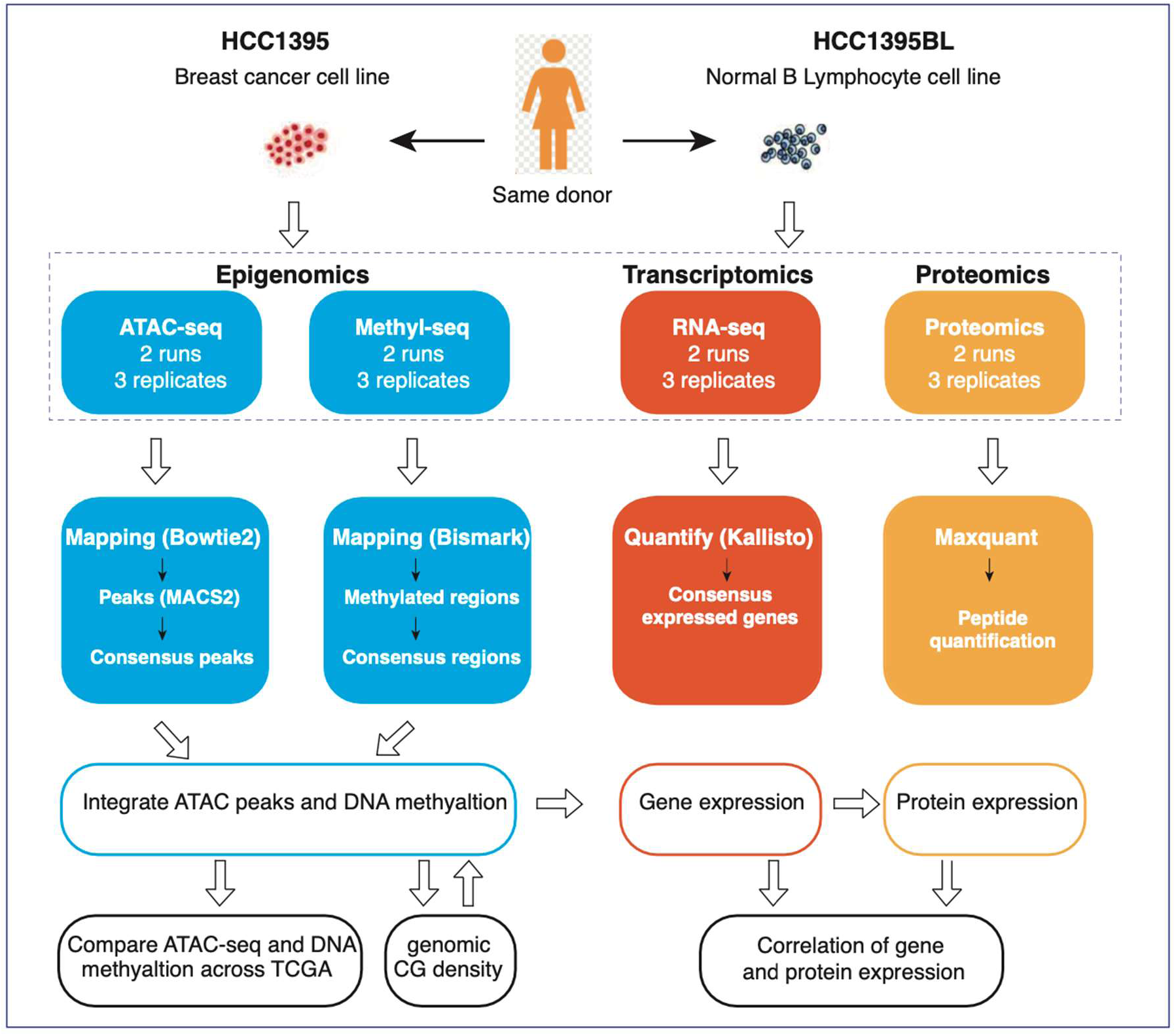
Study design and bioinformatics workflow. HCC1395 and HCC1395BL represent a breast cancer cell line and “normal” (i.e., immortalized) B lymphocyte cell line, respectively. ATAC-seq, RRBS, RNA-seq and proteomic analyses were performed on three replicates of both cell lines. Each data type was analyzed using standard analytical pipelines. ATAC-seq and DNA methylation were analyzed to understand correlation between two layers of epigenetics in relation to genomic CG density. Gene expression was correlated with epigenetic marks and protein expression.

### Annotation of HCC1395 and HCC1395BL ATAC peaks revealed cell line-specific open chromatin

To identify open chromatin regions, we mapped ATAC-seq data to the human genome (hg38) with bowtie2^21^ and used MACS2^16^ to identify peaks across all samples. Visualization of the ATAC peaks revealed that some peaks were detected across all replicates in both cell lines (peaks highlighted in black rectangular box) while others were detected in cell line-specific manner (peaks highlighted in red and blue rectangular boxes) (**Fig. 2a**). We detected approximately around 200,000 ATAC peaks on each replicate (**Fig. 2b**). Correlation of peaks between replicates showed high reproducibility in HCC1395 (R=0.914) and HCC1395BL (R=0.917) (**Fig. S1a**). From the initial pool of peaks, we retained only those peaks that were detected in all three replicates, resulting in 134,320 consensus peaks in HCC1395 and 139,179 consensus peaks in HCC1395BL (**Fig. 2b**). Approximately, one third (48,963) of the consensus (or concordant) peaks were common between the two cell lines (**Fig. 2c**), indicating that majority of the open chromatin regions were specific to each cell line. This finding is consistent with previous observations of a low overlap of ATAC peaks across various tumor types^22^. About 18% of the peaks overlapped promoter regions, while majority (>50%) of the peaks were in gene bodies (intragenic regions) (**Fig. 2d**) (**Supplementary Table 2-3**). Only about 12% of ATAC peaks overlapped with CpG islands (CGIs; regions with high CG density) (**Fig. 2e**).

**Figure 2.**
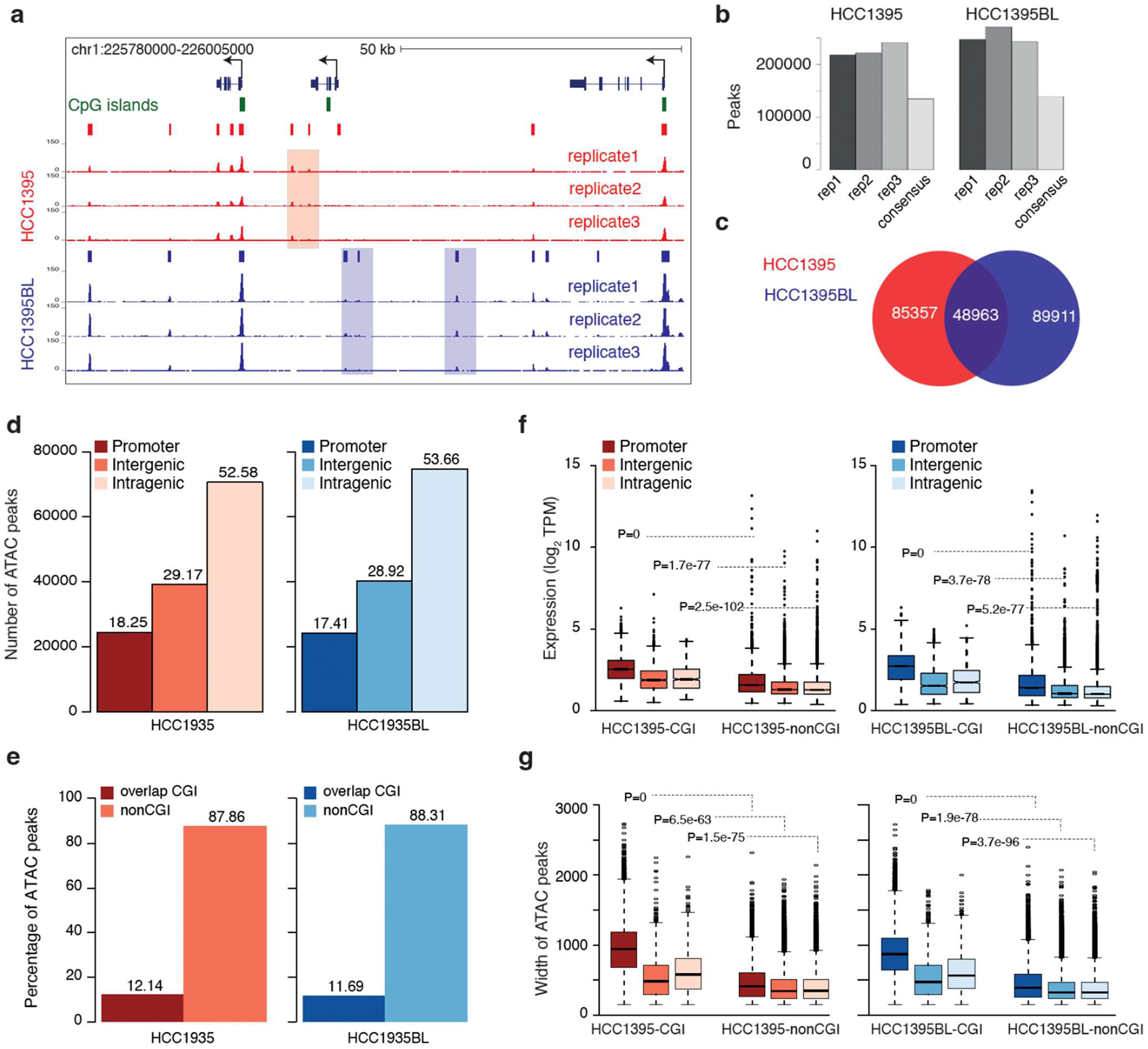
Open chromatin landscape of HCC1395 and HCC1395BL cell lines. **(a)** A UCSC browser view showing 3 replicates of ATAC-seq coverage of HCC1395 (red) and HCC1395BL (blue) cell lines. ATAC peaks region are shown as red and blue rectangular bars. **(b)** Bar plot shows the number of ATAC peaks from three replicates. Peaks identified in all three replicates are defined as consensus peaks. **(c)** Venn diagram shows the overlap of ATAC peaks from the HCC1395 and HCC1395 cell lines. **(d)** Distribution of ATAC peaks with respect to promoter, intragenic, and intergenic regions. **(e)** Distribution of ATAC peaks with respect to CpG islands (CGI). **(f)** Mean counts of ATAC peaks overlapping promoter, intragenic, and intergenic regions. Peaks from each region are further classified based on overlap with CpG islands (CGIs). P-values were calculated using t-tests by comparing CGI and nonCGI groups across promoter, intragenic, and intergenic regions. **(g)** Box plots show the distribution of ATAC peak width across promoter, intragenic, and intergenic regions, which were classified into two groups based on overlap with CGIs.

Thus, most open chromatin regions in both cell lines were in intragenic regions and had low CG density. CGI ATAC peaks had significantly higher mean read count (reflecting higher peak intensities) than in non-CGI ATAC peaks in both cell lines, irrespective of their location in promoter, intragenic, or intergenic regions (**Fig. 2f**). ATAC peaks overlapping promoter CGIs had significantly higher peak intensity than those of intergenic and intragenic CGI ATAC peaks. Similarly, among the non-CGI ATAC peaks, those overlapping promoters had significantly higher intensity than those in both intergenic and intragenic regions (**Fig. 2f**). We also observed that CGI ATAC peaks were significantly broader than non-CGI peaks (**Fig. 2g**; **Fig S1a**) reflecting wider open chromatin regions compared with non-CGI ATAC peaks. Thus, despite there being only a minimal overlap in ATAC peaks between the two cell lines, the genomic context of these peaks in terms of signal intensity and width showed a similar CG-dependent variation, reflecting the influence of genomic CG density influence open chromatin landscape in cell type-specific manner.

### DNA methylation landscapes revealed influence of genomic CG density and chromatin context in cell line-specific methylation patterns

To annotate the DNA methylation landscape of these two cell lines, we used Bismark^17^ to analyze methyl-seq data and identify methylation levels of CGs (mCG). We retained only those CGs whose methylation levels were measured in all three replicates (**Methods**) for each cell line, resulting in 1,314,941 mCGs in HCC1395 and 1,415,665 mCGs in HCC1395BL (**Fig. S2a; Supplementary Table 4-5)**. Around 50% of the detected mCG sites overlapped between the two cell lines (**Fig. S2a**), indicating a significant portion of CpG sites are methylated in a cell type-specific manner. The methylation levels of CpG sites (measured as the *beta* value (see Methods)) revealed a bimodal distribution in both cell lines (**Fig. 3a**), reflecting that most CpGs are either methylated (*beta* value closer to 1) or unmethylated (*beta* value closer to 0), similar to previous observations across human tissues^23^. The average methylation levels across gene bodies revealed low methylation at gene transcription start sites (TSSs) reflecting that the promoter regions were generally unmethylated. Methylation levels progressively increased away from TSSs and peaked towards transcription end sites (TESs). The methylation levels at promoters overlapping CGIs were much lower compared with non-CGI promoters (**Fig. 3b**). Relatively higher methylation levels of non-CGI promoters suggests that some CpGs remain methylated, while almost all CpGs are unmethylated in CGI promoters. This suggests that methylation levels at promoters are tied to genomic CG density.

**Figure 3.**
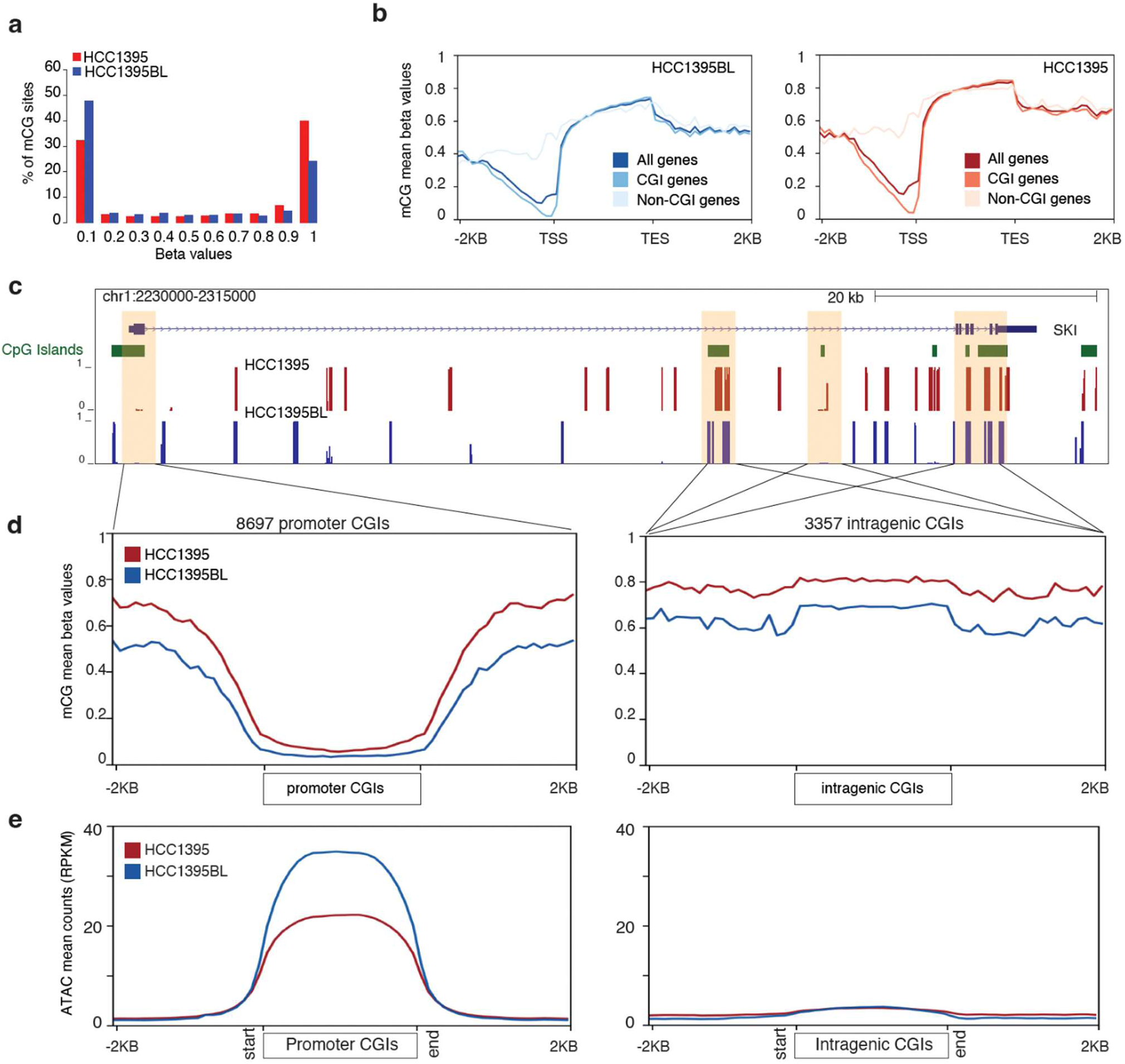
DNA methylation landscape of HCC1395 and HCC1395BL. **(a)** Distribution of beta value of CG methylation. Beta value ranged between 0-1 and are divided into 10 bins. **(b)** Mean methylation levels of all expressed genes. Gene length is scaled between transcription start sites (TSS) and transcription end site (TES). **(c)** A UCSC browser view showing the gene SKI and CpG islands (CGIs) overlapping promoter and intragenic regions. Mean methylation levels of HCC1395 and HCC1395BL are represented as beta values in the range of 0-1. CGIs overlapping promoters have low methylation levels while four intragenic CGIs have high methylation levels. **(d)** Mean methylation levels of promoter CGIs (left panel) and intragenic CGIs (right panel) and 2 KB flanking CGIs. The Y axis represents the mean methylation level (beta value). **(e)** ATAC signals in promoter CGIs (left panel), intragenic CGIs (right panel), and 2 KB flanking regions. The Y axis represents the mean ATAC signal measured in RPKM.

In addition to promoter CGIs, thousands of genes have intragenic CGIs, as exemplified for gene *SKI* (**Fig. 3c**), thus we sought to understand their methylation patterns. Visual inspection revealed that promoter CGI had low methylation while intragenic CGIs were highly methylated (**Fig. 3c**). The methylation levels of all active promoter CGIs (N=8,697) were low across the entire CGI (average promoter CGI length: 2KB) but gradually increased away from CGIs (**Fig. 3d**). In contrast, intragenic CGIs (N=3,357) had high methylation levels than in the flanking gene body^24^ (**Fig. 3d**). Opposite methylation levels between promoter and intragenic CGIs, despite having similar CG density, suggest additional factors might influence methylation levels. To this end, we assessed how the chromatin accessibility was correlated with methylation levels. We observed high intensity of ATAC signals, reflecting open chromatin near promoter CGIs. In contrast, intragenic CGIs had low intensity ATAC signals, reflecting a closed chromatin state (**Fig. 3e**). This indicated that the DNA methylation levels of genomic regions were associated with genomic CG density and its chromatin context.

To understand the association between open chromatin and DNA methylation, we classified ATAC peaks into two groups based on overlap with CGIs. In both groups, ATAC peaks were further classified into four bins based on their CG density. Open chromatin regions with higher CG density had higher ATAC signals in both cell lines (**Fig 4a-b**; left panels). We observed that ATAC signals in open chromatin regions decreased with CG density in both CGI and non-CGI groups (**Fig. 4a-b**; right panels). However, overall ATAC signals were higher in CGI regions. This suggests that chromatin accessibility (as measured by ATAC signals) is largely influenced by genomic CG density. We then plotted methylation levels and observed non-CGI ATAC peaks had higher methylation levels compared to CGI ATAC peaks (**Fig. 4c-d**). While both CGI and non-CGI ATAC peaks represented accessible chromatin regions, relatively high methylation levels in non-CGI peaks revealed some CGs remained methylated despite being in an open chromatin region. This highlights that the association between chromatin accessibility and DNA methylation is influenced by the genomic CG density.

**Figure 4.**
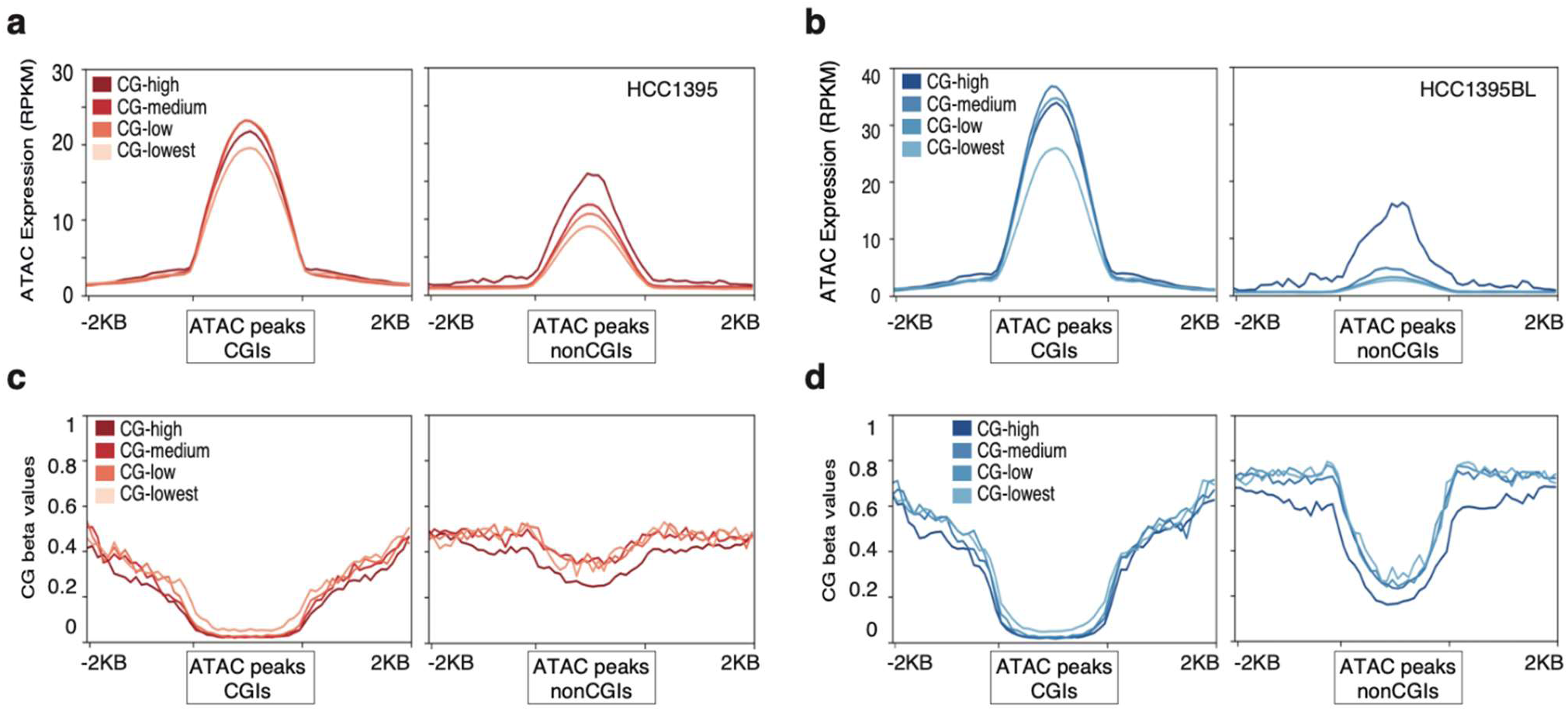
Association of CG density, open chromatin, and CG methylation. **(a)** ATAC peaks overlapping CpG islands (CGIs) (left panel) and non-overlapping CGIs (right panel) in the HCC1395 cell line. ATAC peaks are grouped into 4 bins based on decreasing CG density. **(b)** Same as in A, but for HCC1395 cell line. **(c-d)** CG methylation levels of ATAC peaks overlapping CpG islands (CGIs) (left panel) and non-overlapping CGIs (right panel) in HCC1395 **(c)** and HCC1395BL **(d).** ATAC peaks are grouped into 4 bins based on decreasing CG density. The Y axis indicates methylation level (Beta values) which range between 0-1.

### Transcriptomic landscape and its association with chromatin accessibility and DNA methylation

We next sought to quantify the gene expression levels of reference and alternative transcripts across two cell lines by RNA-seq and understand the association with their promoter epigenetic profiles. We used Kallisto^18^ to quantify the expression levels of all Ensembl transcripts and included only those transcripts that had a minimum of 0.5 tags per million (TPM) in all three replicates and a minimum of 1 TPM in at least one replicate. A total of 18,368 and 19,197 transcripts were detected in HCC1395 and HCC1395BL (**Supplementary Table 6-7**), respectively, representing 10,400 and 10,199 genes (**Fig. 5a**). Around 30% of genes had more than one transcript; these were classified as either reference or alternative transcripts (Methods). We overlapped promoters with CGIs and observed that promoters of reference transcripts in both cell lines had significantly higher overlap with CGIs (**Fig. 5a**), thus reflecting an enrichment of nonCGI promoters among alternative transcripts. Expression levels of reference transcripts were also higher than alternative transcripts in both CGI and nonCGI groups (**Fig. 5b**). Within reference transcripts, promoters overlapping with CGIs had higher expression than nonCGI promoters, indicating that genes with CGIs on promoters had higher expression levels. Overall, lower expression levels of alternative transcripts can partially be explained by their low GC-content.

**Figure 5.**
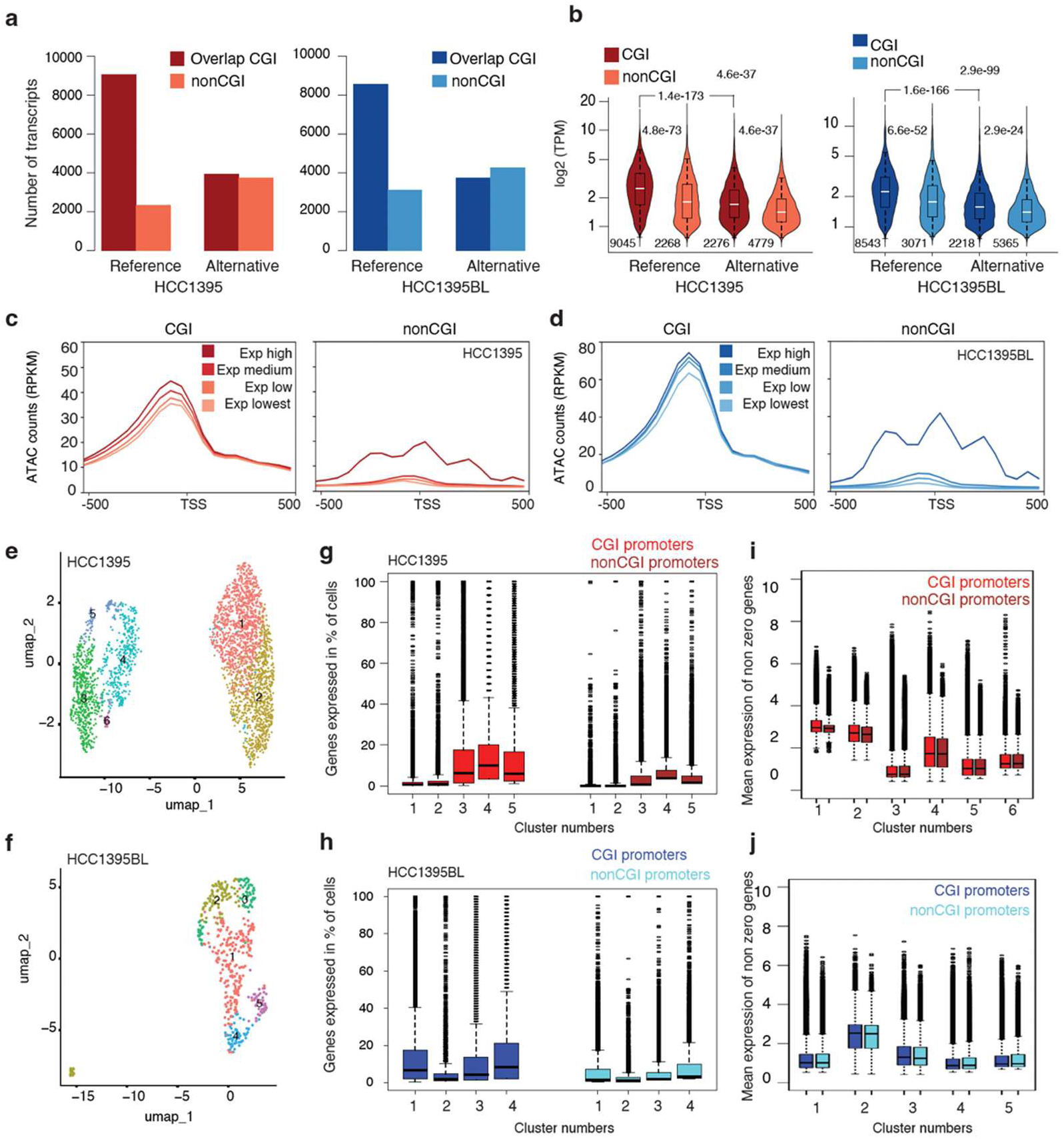
Gene expression profile of the HCC1395 and HCC1395BL cell lines. **(a)** Number of reference and alternative transcripts expressed in HCC1395 (left panel) and HCC1395BL (right panel). Alternative promoters were depleted for CpG islands (CGIs). **(b)** Violin plots show mean expression levels across three replicates for HCC1395 (left panel) and HCC1395BL (right panel). P-values were computed using two-sided t-test. **(c)** The coverage of ATAC-seq reads around genes transcription start site (TSS). Genes are classified into four bins based on expression levels separately for CGI promoters and nonCGI promoters. **(d)** Same as in E, for HCC1395BL cell line. **(e-f)** UMAP clusters of HCC1395 and HCC1395BL **(f).** (**g-h)** Percentage of single cells in clusters that express CGI promoter genes (on left) and nonCGI promoter genes (on right). Boxplots show that CGI promoter genes were expressed in a greater number of cells than nonCGI promoter genes in the same cluster. **(i-j)** Average expression levels CGI promoters and nonCGI promoters’ genes by excluding genes in single cells with zero counts. The mean expression level of CGI promoters was higher than for nonCGI promoters.

To further understand the association of RNA gene expression with ATAC signals at promoters, genes were classified into four bins based on expression levels. Genes with higher expression levels had higher ATAC signals (**Fig. 5c-d**) at both CGI and non-CGI promoters. While nonCGI promoters had an overall lower ATAC signals, such that even the highly-expressed bin from nonCGI promoters had low levels of ATAC signal intensity. We next compared the expression levels of CGI and non-CGI promoters across different bins and observed that the highly expressed bin from non-CGI promoters had higher expression levels compared with the bin with lowest expression of CGI promoters (**Fig. S3a-b**), while it had lower ATAC signals (**Fig. 5c-d**). This suggested that promoter CG density influenced chromatin accessibility and supported the notion that gene expression levels were tightly correlated with the CG content. We next asked how DNA methylation facilitated alternative transcript expression despite high DNA methylation levels across gene bodies. We compared DNA methylation levels between TSSs of reference and alternative transcripts and observed methylation levels decreased at alternative TSSs (**Fig. S3c**) implying a relationship between DNA demethylation and transcription of alternative transcripts promoters.

We next asked whether the observed lower expression of non-CGI promoters in bulk RNA-seq is associated with lower accessibility leading to expression in relatively fewer cells or to an inherent property of genes. To this end, we analyzed scRNA-seq data^8^ of HCC1395 and HCC1395BL and counted the frequency at which CGI and non-CGI promoter genes were expressed in a given single cell across each of the UMAP clusters (**Fig. 5e-f**). In both cell lines, CGI promoter genes were expressed in more cells (**Fig. 5g-h**), which was consistent with high accessibility of these genes. On the other hand, non-CGI promoter genes were expressed in relatively fewer cells (**Fig. 5g-h**). As non-CGI promoter genes were expressed in relatively fewer cells, their average expression levels are expected to be lower in bulk RNA-seq, which is consistent with our observation (**Fig 5b**). Thus, we asked if we should only analyze the cells where nonCGI promoter genes are expressed, how the expression level would compare with respect to CGI promoters. We observed that the expression levels of CGI promoter genes were higher than non-CGI promoters’ genes even when analyzing at single cell level where genes were expressed, (**Fig. 5 i-j**). This highlights that gene expression is influenced by genomic CG density, which in turn modulates epigenetic regulation and gene expression.

### Non-CGI accessible chromatin revealed a tissue-specific regulation across TCGA cancers

While the overlapping of the ATAC peak regions between the two cell lines was low, we sought to understand the variation in overlap of peaks across different tumor types in CG-dependent manner. We then asked whether the association of open chromatin with CpG density, observed in the two reference cell lines, was conserved across 23 cancer types from The Cancer Genome Atlas (TCGA)^22^. The ATAC peaks overlapping CGIs were mostly (>80%) accessible across different tumor tissues (**Fig. 6a**), implying that CGI ATAC peaks are ubiquitously open across different tumor tissues. On the other hand, only 30-50% of the non-CGI ATAC peaks were accessible across other tumor tissues, suggesting that non-CGI ATAC peaks reflected tumor tissue/cell-specific accessible regions. This pattern was consistent in both HCC1395 (**Fig. 6a**) and HCC1395BL (**Fig. 6b**), indicating that despite the low overlap between the two cell lines, CGI and non-CGI ATAC peaks exhibited cell type restricted or ubiquitous accessibility, respectively. Based on the chromatin accessibility and DNA methylation of these two cell lines, we propose a distinct mode of regulation in CGI and non-CGI promoters. The CGI promoters have larger open chromatin regions/peaks which are devoid of DNA methylation, associated with relatively high gene expression. On the other hand, non-CGI promoters have narrow open chromatin regions and contain some levels of methylated CGs within ATAC peaks, which could explain the relatively lower tissue-specific gene expression (**Fig. 6c**). The CG density gradient in these epigenetic regulations provides a plausible fine tune the gene expression levels in a tissue-specific or ubiquitous manner.

**Figure 6.**
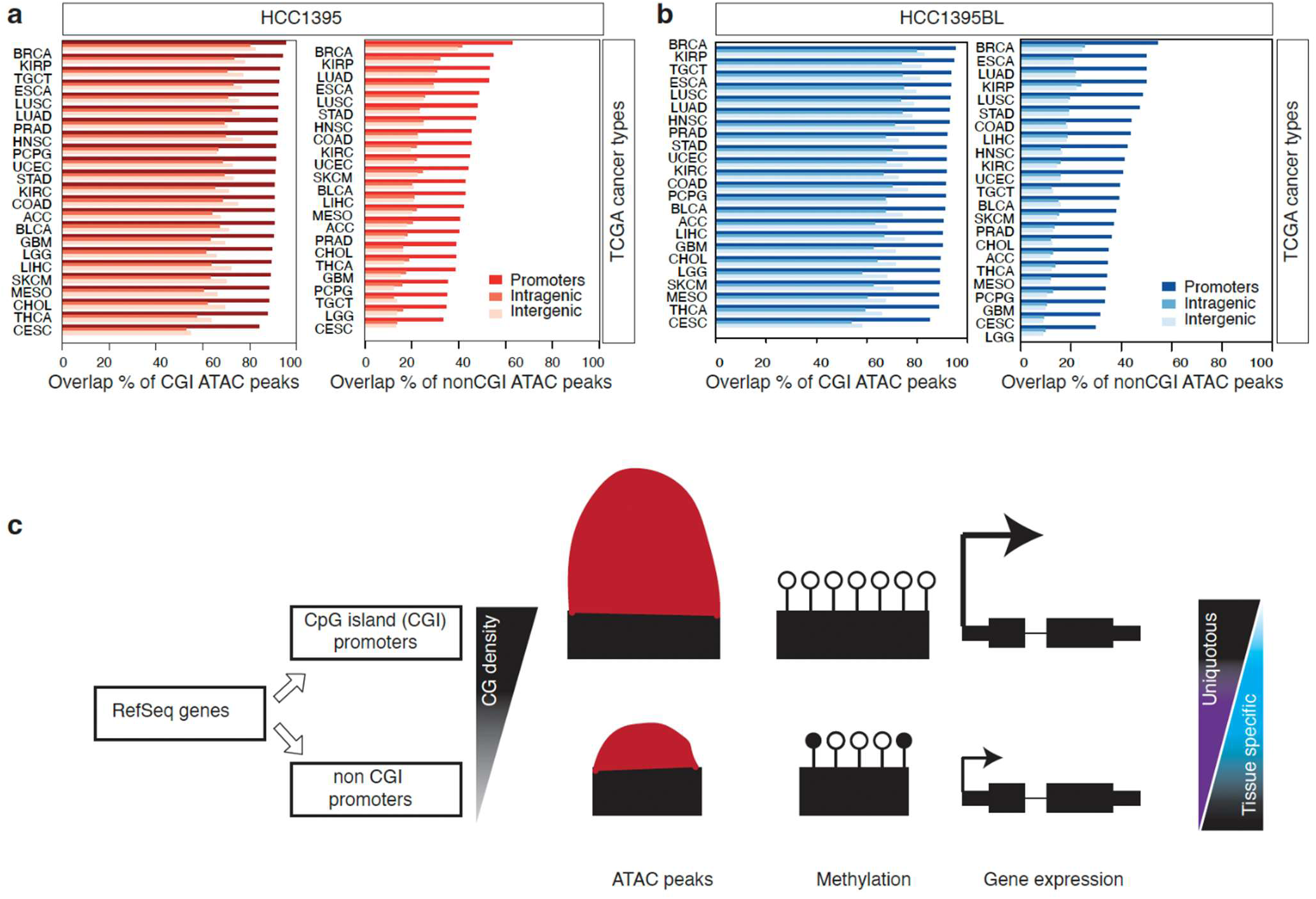
Projection of HCC1395 and HCC1395BL ATAC peaks across 23 tumor tissues from TCGA. **(a)**Overlap of HCC1395 cell line ATAC peaks with the ATAC peaks from 23 different tumor tissues across The Cancer Genome Atlas (TCGA). The majority of HCC1395 CpG islands (CGIs) ATAC peaks (left panel) have detected open chromatin in other tissue types. HCC1395 nonCGI ATAC peaks (right panel) are detected at low levels in other tissue types. X axis indicates the percentage of overlap of HCC1395 ATAC peaks with ATAC peaks from different tumor tissues. **(b)** Same as in a, but for HCC1395BL cell line. **(c)** Schematic representation to show the influence of genomic CG density on epigenetic features (ATAC peaks and DNA methylation) and gene expression.

### Proteomic maps of cell lines and their correlations with the transcriptomes

We next sought to map the protein expression landscape of these cell lines, detect alternatively spliced peptide isoforms, and understand the impact of cell line specific SNVs on protein quantification. We used mass spectrometry and obtained a high coverage of MS/MS spectra (called peptide-spectrum matches) that mapped to multiple regions of peptides as shown for the gene *PFN2* (**Fig. 7a**). Each individual MS/MS spectrum that matched peptides mapping to a particular protein was quantified using MaxQuant^25^. These spectra were then analyzed to quantify mean protein expression levels, including different protein isoforms. For example, the *PFN2* gene encodes for the Profilin-2 protein and has two annotated isoform entries (P35080-1 and P35080-2) in UniProt. Both isoforms are 140 amino acid residues in length and differ in amino acid residues 109 – 140 (**Fig. S4a**). Two isoforms have a different third exon which was supported by mass spectrometry results, where 1 peptide mapped to exon 1 (**Fig. S4b**) and 4 peptides mapped to exon 2 (**Fig. S4c**), common to both isoforms. The third exon had 2 peptides unique to P35080-1 and 3 peptides unique to P35080-2 (**Fig. S4d-e**). Both isoforms also had a unique peptide (VLVFVMGK for P35080-1 and ALVIVMGK for P35080-2) that spanned the splice junction between exon 2 and exon 3 (**Fig. 7a**, **Fig. S4d-e**), supporting the alternative splicing of these isoforms at the protein level.

**Figure 7.**
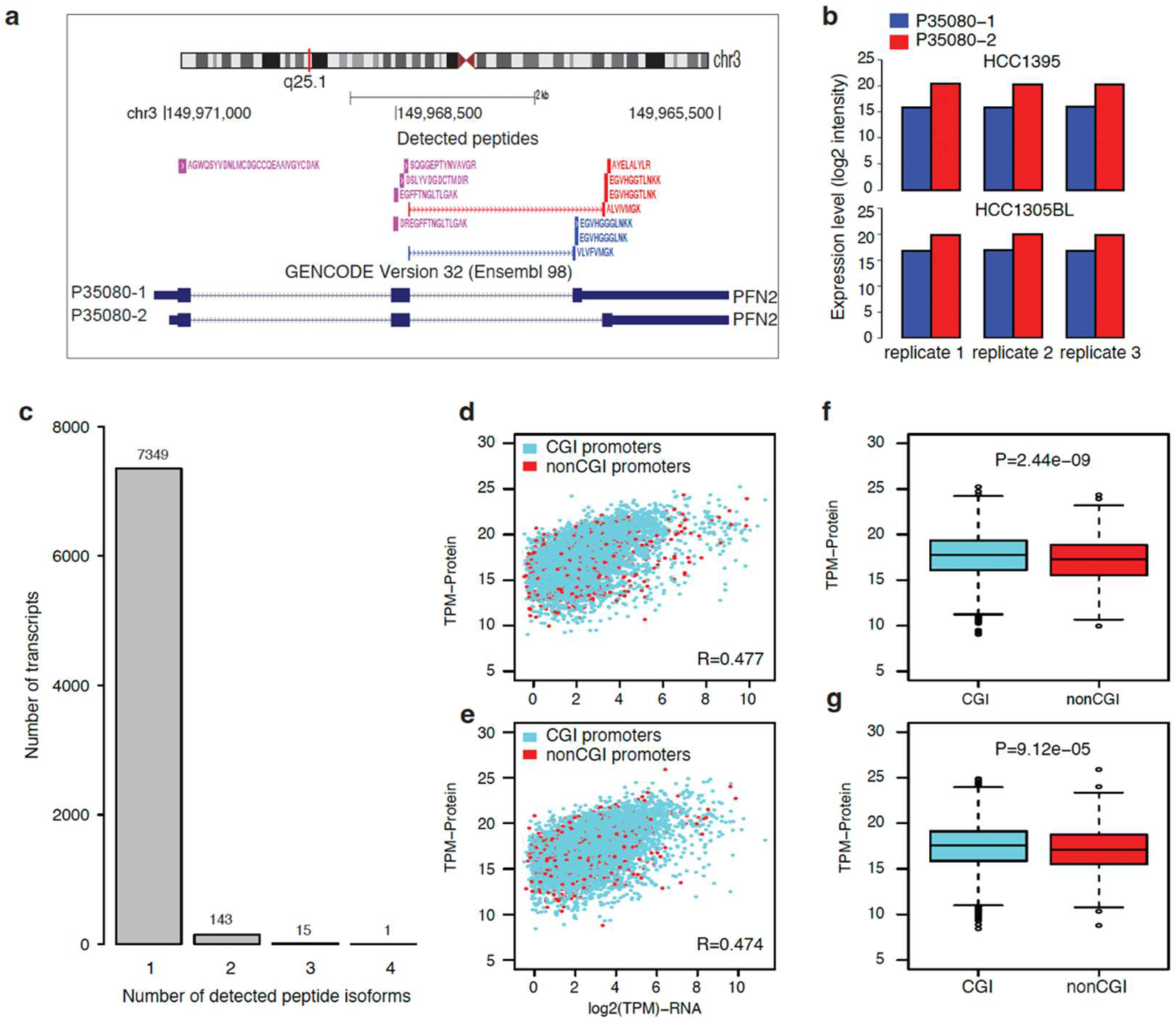
Proteomics map of HCC1395 and HCC1305BL cell lines. **(a)** A UCSC genome browser view (reverse strand) showing peptides identified by MS/MS spectra that mapped to two UniProt isoforms of *PFN2* (P35080-1 and P35080-2). These isoforms share exons 1 and 2 but have different third exons resulting from alternative splicing. Peptides common to both isoforms are colored magenta. Peptides unique to P35080-1 and P35080-2 are colored blue and red respectively. Peptides overlapping in each transcript are summed up to quantify protein expression levels. (**b**) Bar plots show individual protein expression levels of each spliced isoform across three replicates for both HCC1395 and HCC1305BL cells lines. (**c**) Number of peptide isoforms detected per gene. The majority (98.5%) of genes have only one detected peptide. (**d-e**) Correlation of gene expression and protein expression levels. Gene expression and protein levels are positively correlated in HCC1395 (R=0.477) and HCC1395BL (R=0.474). (**f-g**) Average protein expression levels of genes grouped based on overlap with CpG islands (CGI) in their promoters. Genes overlapping CGI have significantly higher protein expression levels in HCC1395 (F) and HCC1395BL (G). P-value was calculated using a t-test.

In total we identified 7,733 peptide isoforms, where the majority (7,349; 95%) of the genes had a single peptide isoform (**Fig. 7c**). We correlated the protein and RNA gene expression levels based on their promoter overlap with CGIs and observed a positive correlation (R=0.477 in HCC1395 and R=0.474 in HCC1395BL) (**Fig. 7d-e**) (**Supplementary Table 8**). The protein expression levels of transcripts overlapping CGIs were significantly higher than protein expression levels of non-CGI transcripts (**Fig. 7f-g**) in both cell lines. This revealed that CGI promoter genes had higher gene and protein expression levels compared with non-CGI promoter genes. Differential expression of protein levels between the two cells lines revealed HCC1395BL to have 614 differentially upregulated proteins (red circles) and 372 significantly downregulated proteins (blue circles) (**Fig. S4f**) at the defined threshold (p-value < 0.05 and absolute log2 ratio > 1).

To understand the effects of somatic mutations on protein expression, we compiled a custom protein FASTA database incorporating somatic SNVs unique to HCC1395 compared with HCC1395BL as identified by the SEQC-2 consortium^1^ (see Methods). We identified a total of 6 variant peptides indicating expression at the protein level of 3 somatic mutations from the truth set in the following genes (*OSTC*, *SEC22B*, and *PRDX5*). In addition, 3 somatic mutations from the non-truth set were observed in *DDX3X*, *FLNA*, and *TUBB8B* (**Supplementary Table 9**). Five of the 6 variant peptides had higher expression in the breast cancer samples compared with the B lymphocyte samples (including all 3 from the truth-set). In addition, four of these amino acid substitutions detected at the protein level were also found in COSMIC from breast ductal carcinoma tissue samples, indicating possible biological relevance: F to L in *OSTC*, F to L in *PRDX5*, R to T in *DDX3X*, and D to H in *FLNA*. To relate the peptides containing the amino acid substitution at the protein level to a specific SNV, we located the specific SNV corresponding to the amino acid substitutions identified by MS-MS. The mapping of the six variant peptides were visualized on genome browser that included three from the truth set (**Fig. S5a-c)** and three from non-truth set (**Fig. S5d-f**). Collectively, we have provided protein expression levels of these two cell lines, association of expression level with genomic CG density and finally showed how often SNVs are incorporated into mutated peptides.

## Discussion

As a follow up to our previous studies^1–3,6–8,15^, here we further provided a comprehensive catalog and detail characterizations at epigenomic, transcriptomic and proteomic levels on two cell line reference samples. We interrogated thousands of open chromatin regions, millions of CG and CH sites on DNA methylation status, thousands of genes on their transcript and protein gene expressions. In addition to the availability of processed datasets, we also demonstrated a comprehensive integrative analysis across different molecular layers of the omic data from the same cell lines. Together with our previous multi-center studies^1–3,6–8,15^ on these two cell lines, we provided well-characterized genomic materials, reference datasets, and reference bioinformatics methods, which will be a valuable resource not only to the cancer research community but also to the genomics research community. From a benchmarking perspective, these cell lines have been annotated with somatic alterations^1–3^ and structural variations^7^, which can provide an opportunity to measure how the personalized somatic alteration will alter epigenetic peak calling. On the other hand, the breast cancer research community can further mine the data to extract novel biological functions associated with cancer.

Our results highlight an overall inverse relationship between chromatin accessibility and DNA methylation. While methylated DNA represents the repressive state, accessible chromatin marks the opening of chromatin of the genomic DNA for active transcription; suggesting that the inverse relationship is biologically relevant. However, we showed that this relationship is dependent upon three factors: 1) transcriptional state (active or inactive); 2) genomic sequence CG density (CG-rich or CG-poor); and 3) genomic regions (promoter, gene body, and intergenic regions). Epigenetic marks of actively transcribed promoters are expected to be different from those of inactive/untranscribed promoters. Thus, when transcriptional states are controlled, genomic CG density largely influences the epigenetic landscapes where CG-rich regions have higher accessibility and lower methylation compared with CG-poor regions (lower accessibility and higher methylation). CG-rich regions not only had higher accessible chromatin and lower methylation levels in a single cell type but showed similar patterns across other cell types. On the other hand, CG-poor regions had restricted chromatin accessibility, which varied across cell types (**Fig. 6a-b**), giving rise to a tissue specific activity. As inverse correlation between chromatin accessibility and DNA methylation is context (genomic CG density, transcriptional state, and genomic regions) dependent, we suggest that future studies consider the context in analyzing and interpreting epigenetics data.

Non-CGI promoters have lower gene and protein expression levels along with cell type-restricted open chromatin (**Fig. 6a-b**). We speculate that the promoter genomic CG density influences epigenetic patterns, which in turn influences the gene and protein. Lower CG density of non-CGI promoters have narrow open chromatin regions, which have certain implications on their gene expressions. As non-CGI promoters’ have narrow open chromatin regions at the promoters, they will have fewer available transcription factor (TF) binding sites. A lower frequency of transcription factor binding sites provides an explanation for lower expression levels. On the other hand, it also suggests that genes might be activated in certain specific contexts and hence explain why they might have cell type-specific expression levels. The overall low expression levels of non-CGI promoters can be explained by CG density, which could in turn regulate epigenetic turnover, and ultimately the expression patterns observed in single cell analyses. Secondly, many CpGs within the open chromatin regions of non-CGI promoters were still methylated (**Fig. 4c-d**), which makes some TF sites inaccessible. As DNA methylation is generally repressive in nature, partially methylated CpGs in promoter regions may explain overall lower expression levels. Thus, lower frequency of TFs and partially methylated CpGs might explain their lower expression levels and tissue restricted expression.

Like epigenomic and transcriptomic profile, the proteomic expression landscape was also correlated with CG density, where CGI promoters also had higher protein expression levels. In addition to quantification of peptide, we also analyzed what fraction of cell line genomic variant are expressed. We identified only 6 SNVs at the protein level, thus confirming some of the mutations are indeed converted to altered peptide. Low detection of SNVs at the protein level suggesting two potential scenarios. Firstly, some of the proteins containing amino acid substitutions might result in unstable proteins that are degraded and remain undetected by mass spectrometry. Secondly, low sequence coverage of certain proteins by mass spectrometry might prevent a peptide containing the variant amino acid substitution from being detected. Of interest, our ability to detect mutations from the non-truth set at the protein level may serve as a method to validate these less confident SNVs, which should also aid in other efforts to benchmark variant and reference calls^26^.

In summary, our study has provided a comprehensive reference epigenomic, transcriptomic and proteomic map of two cell lines studied by SEQC-2. We further demonstrated an integrative approach to analyze epigenomic, transcriptomic and proteomic data to extract meaningful biological observation. Finally, we anticipate that the provided different molecular layers of omics data along with the detailed characterizations with integrative analyses on the two cell lines should serve as a good reference for future use.

## Methods

### ATAC-seq library preparation

ATAC-seq was performed as previously described^12^. Briefly, approximately 50,000 cells per sample were centrifuged at 500 *g* for 5 min in a pre-chilled (4 °C) fixed-angle centrifuge. After centrifugation, cell pellets were then resuspended in 50 µl of cold lysis buffer (10mM Tris-Cl, pH 7.4, 10 mM NaCl, 3 mM MgCl_2_, and 0.1% IGEPAL CA-630) by pipetting up and down three times. This cell lysis reaction was incubated on ice for 3 min. After lysis, 1 ml of ATAC-seq RSB containing 0.1% Tween-20 (without NP40 or digitonin) was added, and the tubes were inverted to mix. Nuclei were then centrifuged for 10 min at 500 *g* in a pre-chilled (4 °C) fixed-angle centrifuge. All supernatants were aspirated and cell pellets were resuspended in 50 µl of transposition mixture. Reactions were incubated at 37 °C for 30 minutes in a thermomixer at 1000 RPM. Immediately following transposition, transposed DNA was purified using a Qiagen MinElute PCR purification kit and amplified for 5 cycles using NEBNext 2x MasterMix. 5 µl (10%) of the pre-amplified mixture was used to determine the number of additional cycles needed by qPCR. The remaining PCR reaction was run to the cycle number determined by qPCR. Finally, the amplified library was purified using a Qiagen MinElute PCR Purification kit.

### Bulk RNA-seq library preparation

We isolated mRNA in bulk from HCC1395 and HCC1395BL cells using the miRNeasy Mini kit (QIAGEN, 217004), and built sequencing libraries using the NuGEN Ovation universal RNA-seq kit. Briefly, 100 ng of total RNA was reverse transcribed and then made into double-stranded cDNA (ds-cDNA) by the addition of a DNA polymerase. The ds-cDNA was fragmented to ∼200 bps using the Covaris S220, and then underwent end repair to blunt the ends followed by barcoded adapter ligation. The remainder of the library preparation followed the manufacturer’s protocol.

### TruSeq methyl capture EPIC library preparation and validation

TruSeq methyl capture libraries were prepared manually following the manufacture’s protocol (document 1000000001643 v01, Illumina). Briefly, cell line DNA samples were quantified using the Qubit high sensitivity double-stranded DNA assay (Invitrogen). 500 ng of DNA was normalized to 52 *µ*l of resuspension buffer (Illumina) and was sheared by sonication with a Covaris S220 to a target size of 160 bp. Illumina Sample Purification Beads (SPB, Illumina reference number, 15037172; manufactured by Beckman coulter) were used at a DNA to beads ratio of 1:1.6 for cleanup and size selection. Insert fragment sizes were verified by a TapeStation 2200 using D1000 Screentape (Agilent Technologies). Double size selection was performed using SPB beads following the end-repair. After adenylating the 3-prime end, index adapters were immediately ligated, and the ligated libraries were purified using SPB beads. Probe hybridization was carried out at 58 °C for 2 hours after an initial denaturing step at 95 °C for 10 minutes. The hybridized probes were then captured using Streptavidin Magnetic Beads (Dynabeads, Invitrogen). After elution, the hybridized probes were subjected to second hybridization, second capture, second elution, and bisulfite conversion with lightning conversion reagent (Zymo Research). After bisulfite conversion, the library was amplified using Kapa HiFi Uracil kit (KaPa Biosystems Pty, Cape Town, South Africa) under the following conditions: 12 cycles of 98 °C for 20 seconds, 60 °C for 30 seconds, 72 °C for 30 seconds; and 72 °C for 5 minutes. The amplified library was purified using SPB beads.

### Library quality control and sequencing

All the libraries were quantified with a TapeStation 2200 (Agilent Technologies) and Qubit 3.0 (Life Technologies). Sequencing was performed on either the NextSeq 550 or HiSeq 4000 platform using SBS reagents.

### Mapping of ATAC-seq data

The ATAC-seq reads were mapped to the human genome (hg38) using Bowtie2^21^. We used MACS to call the peaks individually in all samples. To identify consensus peaks, ATAC peaks had to be present in all three replicates in each cell line. For coverage plots, we generated bigwig files using deepTools^27^.

### Mapping of RNA-seq data

The RNA-seq data were analyzed using Kallisto ^18^. We quantified the GENCODE transcripts (hg38) in TPM using Kallisto. To identify genes/transcripts that are expressed reliably in each cell line, we defined a threshold of 0.1 TPM in all three replicates, while at least one of the replicates was required to have a minimum of 0.5 TPM.

### Mapping of methyl-seq data

The methyl-seq reads were trimmed using trim-galore with command “trim_galore --non_directional –rrbs”. Trimmed reads were mapped to the genome (hg38) using bismark^17^ with the following parameters “bismark --phred33 --bam --non_directional –unmapped”. Methylation calls were extracted using the following parameter “bismark_methylation_extractor bamfile --ignore 2 --ignore_3prime 2 --comprehensive –bedGraph”. To define the consensus call, every methylation had to be detected in all three replicates and have a minimum coverage of 3 reads. Methylation levels for each CG were defined as the ratio of number of detected methylated CpGs to the number of detected methylated and unmethylated CpGs, which ranged between 0-1. This value is the beta value. For visualization of DNA methylation levels across the genome, we built a bigwig coverage track from the beta value. Three replicates of a given cell line were averaged to get a mean methylation value. The mean methylation values were used for the coverage tracks.

### Mapping and analysis of single cell RNA data

The single cell RNA-seq data of HCC1395 and HCC1395BL were downloaded^8^. Single cell RNA data were analyzed using standard Seurat package^19^. For comparing the expression level of genes with CGI and nonCGI promoters, we only compared the expression across cell if the gene was expressed in that cell.

### Classification of CpG islands

We downloaded annotated CpG islands (CGIs) from the UCSC database ^28^. We intersected CGIs with promoter regions of gene transcripts and assigned overlapping CGIs as promoter CGIs. The remaining CGIs were either overlapped with gene body and classified as intragenic CGIs or else classified as intergenic CGIs.

### Observed/expected CG ratio

The ratio of observed/expected (O/E) CG dinucleotides for a given genomic window was calculated using the following formula O/E CG = (CG count) (genomic window) / (C count * G count). We used the "bedtools nuc” function from Bedtools ^29^ to measure these values and computed O/E CG ratio for each genomic window.

### Visualization of metaplots across genes and CGIs

All metaplots across genes and CGIs were plotted using deepTools2 ^27^. From mapped BAM files, we first generated coverage as Bigwig tracks, which are normalized as RPKM using default parameters from deepTools. For the genes and CGIs metaplot, variable length of genes and CGIs were scaled between start and end.

### Proteomics sample preparation

Cell pellets were resuspended in 8 M Urea, 50 mM Tris-HCl pH 8.0, reduced with dithiothreitol (5 mM final) for 30 min at room temperature, and alkylated with iodoacetamide (15 mM final) for 45 min in the dark at room temperature. Samples were diluted 4-fold with 25 mM Tris-HCl pH 8.0, 1 mM CaCl2 and digested with trypsin at 1:100 (w/w, trypsin: protein) overnight at room temperature. Three biological samples were used for each condition, resulting in a total of 9 samples. Peptides were cleaned using homemade C18 stage tips, and the concentration was determined (Peptide assay, Thermo Scientific 23275). 50 µg of each sample was then used for labeling with isobaric stable tandem mass tags (TMT^10^-126, 127N, 127C, 128N, 128C, 129N, 129C, 130N and 131, Thermo Scientific) following the manufacturer’s instructions. The mixture of labeled peptides was fractionated using strong cation exchange beads (Source 15S, GE Healthcare). After the sample was applied, we eluted the beads sequentially with a buffer containing 25% acetonitrile, 0.05% formic acid, and 50, 125, 200, or 400 mM ammonium bicarbonate, respectively. Each SCX fraction was further separated into 8 fractions using a C18 stage tip with a buffer of 10 mM trimethylammonium bicarbonate (TMAB), pH 8.5 containing 5 to 50% acetonitrile. A couple of fractions with low peptide content where combined, resulting in a total of 28 fractions.

### Mass spectrometry and proteomics

Dried peptides were dissolved in 0.1% formic acid, 2% acetonitrile in water. 0.5 μg of peptides from each fraction were analyzed on a Q-Exactive HF-X coupled with an Easy nanoLC 1200 (Thermo Fisher Scientific, San Jose, CA). Peptides were loaded on to a nanoEase MZ HSS T3 Column (100Å, 1.8 µm, 75 µm x 250 mm, Waters). Analytical separation of all peptides was achieved with 100-min gradient. A linear gradient of 5 to 10% buffer B over 5 min, 10% to 31% buffer B over 70 min, 31% to 75% buffer B over 15 min was executed at a 300 nl/min flow rate followed by a ramp to 100%B in 1 min and 9-min wash with 100%B, where buffer A was aqueous 0.1% formic acid, and buffer B was 80% acetonitrile and 0.1% formic acid. MS experiments were also carried out in a data-dependent mode with full MS at a resolution of 120,000 followed by high energy collision-activated dissociation-MS/MS of the top 20 most intense ions with a resolution of 45,000 at *m/z* 200. High energy collision-activated dissociation-MS/MS was used to dissociate peptides at a normalized collision energy of 32 eV in the presence of nitrogen bath gas atoms. Dynamic exclusion was 45 seconds.

### Raw proteomics data processing and analysis

Peptide identification and quantification with tandem mass tags (TMT) reporter ions was performed using MaxQuant^25^ software version 1.6.0.16 (Max Planck Institute, Germany). Protein database searches were performed against the UniProt human protein sequence database (UP000005640). A false discovery rate (FDR) for both peptide-spectrum match (PSM) and protein assignment was set at 1%. Search parameters included up to two missed cleavages at Lys/Arg on the sequence, oxidation of methionine, and protein N-terminal acetylation as a dynamic modification. Carbamidomethylation of cysteine residues was considered as a static modification. Peptide identifications were reported by filtering of reverse and contaminant entries and assigning to their leading razor protein. Data processing and statistical analysis were performed on Perseus^30^ (Version 1.6.0.7). Protein quantitation was performed on biological replicates and two-sample t-test statistics were used with a p-value of 5% to report statistically significant protein abundance.

### Proteogenomic mapping of peptides to genomic coordinates

To map peptides identified in peptide spectrum to their genomic location, we constructed a custom FASTA protein database. To the header for each protein sequence entry, we added the genomic mapping information that contained genomic coordinates for the protein sequence, including the start and end, and CDS (coding sequences) start coordinates (relative to start of genome start position for the protein) and CDS lengths. This header information formatting was based on the proteogenomic data integration tool QUILTS^31^. We downloaded protein-coding transcript translation sequences (gencode.v32.pc_translations.fa) and the GFF3 comprehensive gene annotation (gencode.v32.annotation.gff3) from GENCODE^32^ for GENCODE human Release 32 (GRCh38.p13). The genomic mapping information for each protein entry in the FASTA database was obtained from entries in the gene annotation. Knowing the position of the peptide in the protein sequence, and the mapping information of the protein sequence relative to the genome, allowed each peptide in the MaxQuant search to be mapped to the genome. A BED file, containing all the peptides with their mapping information, enabled display of the peptides in UCSC genome browser.

To map peptides to somatic missense amino acid substitutions, we constructed protein FASTA databases containing somatic missense SNVs (SNV.MSDUKT.superSet.v1.1.vcf.gz) from the HCC1395 which was downloaded from https://ftp-trace.ncbi.nlm.nih.gov/seqc/ftp/Somatic_Mutation_WG/release/v1.1/, and produced a list of somatic mutations used to screen our proteomic data. We divided the Variant Call Format (VCF) file into two files: (1) a truth set of somatic mutations and (2) a non-truth set of somatic mutations. The truth set contains VCF entries with a PASS label denoting high-confidence somatic calls, while the non-truth set contains the remaining VCF entries. We ran the Ensembl Variant Effect Predictor^33^ (VEP) tool to identify missense variants in these 2 VCF files. We made separate FASTA databases for the truth- and non-truth sets of missense variants. These FASTA databases were constructed using gene annotations as was done for the reference FASTA database, except that the variant data were used to modify the protein sequence to include missense amino acid changes. The position of the variation was maintained in the FASTA header following the format used in QUILTS. The somatic mutations were annotated using the VEP (Variant Effect Predictor) from Ensembl. We filtered the somatic mutation dataset to include only missense mutations in protein coding regions. We produced 2 FASTA database files: one for true somatic mutations (truth set) with high confidence calls as defined by SEQC-2 and another for the remaining somatic mutations (non-truth set).

We conducted a MaxQuant search using the reference FASTA protein database and the 2 SNV protein databases containing the somatic missense amino acid substitutions. The MaxQuant output file (peptides.txt) lists each peptide identified as a peptide-spectrum match by the search and information related to its associated FASTA header(s) for the protein sequence(s) in which it can be found. With this it was possible to identify both peptides from the reference as well as variant peptides containing missense substitutions. We determined the position of the peptide in the protein sequence and used the mapping information contained in the FASTA header to produce a header track line in browser extensible data (BED) files that maps the peptide to the genome. Some peptides map to more than one genomic location. Each genomic location (chromosome name, start coordinate, stop coordinate, strand, and any sub-blocks for peptides spanning introns) serves as an address for a peptide. A BED file was produced for the mapping location of the peptides, which can be visualized in a genome browser.

## Data availability

The datasets generated and analyzed (multi-center scRNA-seq data) in the current study are available in the SRA with the accession code PRJNA504037 at https://www.ncbi.nlm.nih.gov/bioproject/PRJNA504037. The ATAC-seq data of 23 tumor tissues from TCGA are downloaded from the following URL: https://gdc.cancer.gov/about-data/publications/ATACseq-AWG.

The data of ATAC-seq, Methyl-seq for the two cell lines are available at GEO with the accession code GSE268608 and the URL as the following. The reviewer’s token is available upon request. https://urldefense.com/v3/https://www.ncbi.nlm.nih.gov/geo/query/acc.cgi?acc=GSE268608__;!!PCzdSas!MHdAnRnlJeHaYZsEwk5hOppjNsSitdsNYbsRsARKpsnniN04UHPNAa4Qox6xsS4RIyaRFgP8paz67WDhLw$

The mass spectrometry proteomics data have been deposited to the ProteomeXchange Consortium via the PRIDE^34^ partner repository with the dataset identifier project accession code PXD052353 with the following URL. The reviewer’s token is available upon request. https://www.ebi.ac.uk/pride/review-dataset/922723dc26ce4d5097b3a43c43b4070d

## Code availability

We used many algorithms and code sets for our bioinformatics analyses which have been published previously. All code and software used will be available in Github.

## Acknowledgements

The authors would like to thank Ms. Diana Ho and Ms. Adriana Lopez of the LLU Center for Genomics for their administrative support, particularly in coordinating the Zoom conference calls for the project. The authors would like to thank ATCC, and particularly Liz Kerrigan for providing the two cell lines, i.e., HCC1395 and HCC1395BL for our study. The genomic work carried out at the LLU Center for Genomics was funded in part by the National Institutes of Health (NIH) grants S10OD019960 (CW), U01DA058278 (CW), the Ardmore Institute of Health grant 2150141 (CW) and Dr. Charles A. Sims’ gift to LLU Center for Genomics. The work of Chunlin Xiao was supported by the National Center for Biotechnology Information of the National Library of Medicine (NLM), National Institutes of Health.

## Authors’ contributions

CW conceived, designed the overall study, and provided funding. XC designed and funded the proteomics study. CW managed the project. CN drafted the manuscript and conducted all the bioinformatics data analyses. WC, ZC and WL carried out genomics experiments and helped the data analysis. LX carried out proteomics experiments and JW performed the proteomics data analysis. CX, WJ, MMJ, AF and CW helped edit the manuscript. All authors reviewed the manuscript. CW revised and finalized the manuscript.

## Competing interests

Andrew Farmer is an employee of Takara Bio USA, Inc., and Wendell Jones is an employee of Q^2^ Solution company. All other authors claim there are no conflicts of interest. The views presented in this article do not necessarily reflect the current or future opinion or policy of the US Food and Drug Administration. Any mention of commercial products is for clarification and not intended as an endorsement.

## Notes

### Competing Interest Statement

Andrew Farmer is an employee of Takara Bio USA, Inc., and Wendell Jones is an employee of Q2 Solution company. All other authors claim there are no conflicts of interest. The views presented in this article do not necessarily reflect the current or future opinion or policy of the US Food and Drug Administration. Any mention of commercial products is for clarification and not intended as an endorsement.

### Summary of Updates

1. Removed the reviewer's token to access the original raw data to keep the raw data confidential until the ms is accepted for a publication. 2. Removed the GitHub link for the code to keep it confidential until the ms is accepted for a publication. 3. Updated the author contribution and acknowledgement texts.

